# Rat primary cortical cell tri-culture to study effects of amyloid-beta on microglia function

**DOI:** 10.1101/2024.03.15.584736

**Authors:** Hyehyun Kim, Bryan Le, Noah Goshi, Kan Zhu, Ana Cristina Grodzki, Pamela J. Lein, Min Zhao, Erkin Seker

## Abstract

**Introduction:** The etiology and progression of sporadic Alzheimer’s Disease (AD) have been studied for decades. One proposed mechanism is that amyloid-beta (Aβ) proteins induce neuroinflammation, synapse loss, and neuronal cell death. Microglia play an especially important role in Aβ clearance, and alterations in microglial function due to aging or disease may result in Aβ accumulation and deleterious effects on neuronal function. However, studying these complex factors *in vivo*, where numerous confounding processes exist, is challenging, and until recently, *in vitro* models have not allowed sustained culture of microglia, astrocytes and neurons in the same culture. Here, we employ a tri-culture model of rat primary neurons, astrocytes, and microglia and compare it to co-culture (neurons and astrocytes) and mono-culture enriched for microglia to study microglial function (i.e., motility and Aβ clearance) and proteomic response to exogenous Aβ.

**Methods:** We established cortical co-culture (neurons and astrocytes), tri-culture (neurons, astrocytes, and microglia), and mono-culture (microglia) from perinatal rat pups. On days *in vitro* (DIV) 7 – 14, the cultures were exposed to fluorescently-labeled Aβ (FITC-Aβ) particles for varying durations. Images were analyzed to determine the number of FITC-Aβ particles after specific lengths of exposure. A group of cells were stained for βIII-tubulin, GFAP, and Iba1 for morphological analysis via quantitative fluorescence microscopy. Cytokine profiles from conditioned media were obtained. Live-cell imaging with images acquired every 5 minutes for 4 hours was employed to extract microglia motility parameters (e.g., Euclidean distance, migration speed, directionality ratio).

**Results and discussion:** FITC-Aβ particles were more effectively cleared in the tri-culture compared to the co-culture. This was attributed to microglia engulfing FITC-Aβ particles, as confirmed via epifluorescence and confocal microscopy. Adding FITC-Aβ significantly increased the size of microglia, but had no significant effect on neuronal surface coverage or astrocyte size. Analysis of the cytokine profile upon FITC-Aβ addition revealed a significant increase in proinflammatory cytokines (TNF-α, IL-1α, IL-1β, IL-6) in tri-culture, but not co-culture. In addition, Aβ addition altered microglia motility marked by swarming-like motion with decreased Euclidean distance yet unaltered speed. These results highlight the importance of cell-cell communication in microglia function (e.g., motility and Aβ clearance) and the utility of the tri-culture model to further investigate microglia dysfunction in AD.

## Introduction

Alzheimer’s Disease (AD) is a progressive neurodegenerative brain disorder that impairs memory and cognitive function and is the most common cause of dementia, with an estimated 6.7 million Americans living with the condition (Alzheimer’s Association, 2023). While the pathogenesis of AD is unclear, abnormal deposits of amyloid-β (Aβ) plaques and hyperphosphorylated tau proteins throughout the brain are thought to play a role in neuroinflammation, synapse loss, and neuronal cell death (Selkoe and Hardy, 2016). While many different cell types, most importantly neurons, astrocytes, oligodendrocytes, and microglia, constitute the complex functions of the central nervous system (CNS), microglia (resident macrophages) have attracted considerable interest for their central role in homeostatic and pathological CNS states (Streit et al., 2004). In homeostasis, microglia continuously survey the CNS to engulf cellular and biomolecular debris (Neumann et al., 2009) such as Aβ, support neuronal function via pruning synapses (Schafer et al., 2012), and regulate myelination (Cignarella et al., 2020). However, chronic microglial activation – a prominent hypothesis on the etiology of AD – may produce prolonged secretion of pro-inflammatory cytokines, which in turn can exacerbate Aβ accumulation and neuronal cell loss (Wang et al., 2015). In addition, microglia exhibit changes in motility (e.g., clustering, impairment of directional motility) in inflamed tissue with aberrant Aβ proteins (Franco-Bocanegra et al., 2019). However, it is not clear whether these changes are due to the Aβ particles themselves or are mediated through neuron-astrocyte-microglia crosstalk, where all cell types are exposed to Aβ.

While *in vivo* models provide a holistic approach to the study of Aβ accumulation/clearance and its effects on cognition and behavior, *in vitro* models are better suited to examine the molecular mechanisms underlying the interaction of CNS cells with Aβ particles, including receptor-mediated phagocytosis (Koenigsknecht and Landreth, 2004) and the influence of microglial cytokines (Guedes et al., 2018) . To that end, several studies have established various permutations of neuron-glia cultures (e.g., neuron-astrocyte (Chapman et al., 2016), neuron-microglia (Rueda-Carrasco et al., 2023) using cell lines (De Simone et al., 2017; Peng et al., 2021), primary mouse cells (Rueda-Carrasco et al., 2023), primary rat cells (Sahu et al., 2019), primary porcine cells (Aubid et al., 2019), and human primary cells (Ray et al., 2014). In addition, several studies have employed human induced pluripotent stem cell-based (hiPSC) cultures. While human cells, if successfully differentiated with high yield, which is a laborious process, may have higher biological relevance, these cultures still have high variability, thus reducing experimental reproducibility (Jones et al., 2017; Volpato and Webber, 2020). In addition, it is difficult to reliably and consistently generate large numbers of iPSC-derived neurons that mature to the point of extending distinct axons and dendrites and firing action potentials (Prè et al., 2014) These features dictate neuronal function (e.g., electrophysiological activity), which plays a critical role in the cognitive impairment that is the hallmark clinical phenotype of AD. Rodent cell culture models, while not perfect, are valuable in providing fully-differentiated neurons that exhibit robust electrophysiological activity and still serve as critical tools in neuroinflammation and neurodegeneration research (Nazem et al., 2015; Goshi et al., 2020).

We have recently developed an enhanced primary rat cortical cell culture model comprised of the three major cell types associated with neuroinflammation (i.e., neurons, astrocytes, microglia) that can be sustained in a serum-free media formulation to preserve microglia in their homeostatic phenotype (Goshi et al., 2020). The model more faithfully captured CNS response to lipopolysaccharide (LPS) exposure, glial scarring in a scratch assay, and neuronal rescue from glutamate-induced excitotoxicity. In addition, we have used the tri-culture to establish models of toll-like receptor agonist-induced neuroinflammation (Goshi et al., 2022) and used it to monitor electrophysiological changes due to exposure to LPS (Goshi et al., 2023). Here, we employ the tri-culture to compare Aβ clearance by microglia and assess changes in cytokine levels upon Aβ exposure in tri-culture vs. co-culture (neurons and astrocytes). In addition, we quantify microglia motility within the tri-culture or as a mono-culture to provide insight into the potential role of neuron-astrocyte-microglia crosstalk in mediating microglia motility behavior.

## Materials and Methods

### 2.1 Primary cortical co- and tri-cultures

Rat primary cortical cells were obtained from postnatal (day 0-2) Sprague-Dawley rat pups, as previously described (Wayman et al., 2012; Goshi et al., 2020). All studies were conducted according to protocols approved by the Institutional Animal Care and Use Committee of the University of California, Davis. Prior to seeding the cells, the culture plate were coated with 0.5 mg/mL of poly-L-lysine (PLL) in B-buffer (3.1 mg/mL boric acid and 4.75 mg/mL borax, Sigma) at 37°C and 5% CO_2_ for 4 hours, then washed with sterile deionized water six times before covering with the plating medium (Neurobasal A culture medium, 2% B27 supplement, 1x GlutaMAX, 10% heat-inactivated horse serum, and 20 mM HEPES) overnight. The co-culture medium was composed of Neurobasal A culture medium (Thermo Fisher), 2% B27 supplement (Thermo Fisher), and 1x GlutaMAX (Invitrogen). To create the tri-culture medium, mouse IL-34 (R&D Systems) TGF-β2 (Peprotech), and ovine wool cholesterol (Avanti Polar Lipids, Alabasters) were added to the co-culture medium. IL-34 and TGF-β2 have a limited shelf life, necessitating the preparation of a fresh batch of tri-culture medium on a weekly basis.

### 2.2 Primary microglia mono-culture

Microglia were mechanically isolated from tri-cultures at DIV 7 by gently tapping the tri-culture plate against the surface multiple times until most of the microglial cells detached from the culture surface. The detached cells were suspended in fresh tri-culture media. Subsequently, the cells were seeded onto PLL-treated tissue culture plate at a density of 90 cells per mm^2^. The cultures were kept in a humidified incubator at 37°C with 5% CO_2_ with half-volume media replacements every 3-4 days during the course of the experiment.

### 2.3 FITC-Aβ_1-42_ preparation

To prepare Aβ, 0.5 mg of FITC tagged Aβ_1-42_ peptide (Bachem, #4033502) was initially dissolved in 60 μL DMSO (Thermo Fisher) at room temperature and resuspended in cold Dulbecco’s phosphate-buffered saline solution with calcium and magnesium (DPBS+) to achieve 100 μM final stock concentration. The solution was then vortexed for 30 seconds followed by 5-minute incubation at room temperature, repeating for three times. The peptide solution was aliquoted and stored at -80°C until use. FITC-Aβ_1-42_ peptide were characterized using silver staining and immunoblotting (Figure S1). Prior to cell addition, Aβ peptide solution was vortexed and incubated at 4°C overnight. At DIV 7-10, half of the medium in each well was replaced with 2 times higher concentration of the Aβ. For the vehicle control group sterile 2% DMSO in DPBS+ was added. The cultures were then kept at varying times for different experimental purposes from 5 - 96 hours before being evaluated.

### 2.4 Immunostaining and morphological analysis

The cell cultures were washed with warm DPBS+ three times and fixed using 4% PFA in DPBS+ for 20 minutes, as previously described (Goshi et al., 2020). Fixed cultures were permeabilized with 0.05% v/v Tween 20 followed by 0.1% v/v Triton X-100 in DPBS+ and blocked in 5% v/v goat serum with 0.3 M glycine in DPBS+ for an hour prior to 1-hour incubation of primary antibody solution consisting of mouse anti-βIII-tubulin (1:500, Thermo Fisher, RRID: AB_2536829), rabbit anti-GFAP (1:100, Thermo Fisher, RRID: AB_10980769), and chicken anti-Iba1 (1:500, Synaptic Systems 234 009), all in the blocking buffer. The samples were then incubated for 1 hour with a secondary antibody solution that contained goat anti-mouse conjugated to AlexaFluor 647 (diluted 1:500, Thermo Fisher, RRID: AB_2535804), goat anti-rabbit conjugated to AlexaFluor 488 (1:500, Thermo Fisher, RRID: AB_143165), and goat anti-chicken conjugated to AlexaFluor 555 (1:500, Thermo Fisher, RRID: AB_2535858) in DPBS+. Lastly, the samples were counter-stained with 4’,6-diamidino-2-phenylindole (DAPI) solution (1:20,000 in DPBS+) for 5 minutes. Subsequently, the cells were washed with 0.05% v/v Tween 20 in DPBS+ before imaging or mounting the coverslips onto glass slides using Prolong Gold Antifade Mountant (Thermo Fisher). For clearance experiments using FITC-tagged Aβ, only βIII-tubulin and Iba1 stains were used.

The general morphology of stained cells was acquired using a Zeiss Observer D1 microscope at 100x or 200x magnifications. A Leica TCS SP8 STED 3x confocal microscope (Leica Microsystems) with a 63x/1.4 oil immersion objective was used to obtain z-stack images to confirm the phagocytosis of Aβ by microglia. All images were subsequently processed by custom Fiji/ImageJ (Schindelin et al., 2012) macros for better visualization and background signal reduction. The average percent coverage of neurons was assessed by applying default auto-threshold on βIII-tubulin (neurons) images. The sizes of microglia and astrocytes were determined by manually tracing the cell borders and obtaining the cellular footprint (Goshi et al., 2022).

### 2.5 FITC-Aβ_1-42_ peptide clearance assay

The co-cultures and tri-cultures were plated on PLL-treated polystyrene culture wells or glass cover slips in a 24-well plate and maintained for 10 days. FITC-Aβ solution was added to DIV 10 cultures at a final working concentration of 1 μM, with subsequent incubation for 48 - 96 hours. The cultures were fixed with 4% paraformaldehyde (PFA) in PBS for 20 minutes and immunostained as described above. Coverslips with the fixed cells were mounted to glass microscope slides cell side down, and cultures in wells were imaged directly using an inverted fluorescence microscope (Zeiss Observer D1) at 100x magnification. To quantify real-time clearance of particles, we used a Celloger Mini Plus automated live cell imaging system (CURIOSIS) with 40x magnification to image the cultures every 4 hours for 52 hours inside the incubator at 37°C and 5% CO_2_. Fiji/ImageJ software was used to quantify the particles in the culture. The images were batch-processed by applying an auto-threshold and counting with the “Particle Analysis” function to quantify the number FITC-Aβ particles in each image. Number of particles in co-cultures and tri-cultures was counted and presented as number of particles per frame area of the field of view (FOV). The number of particles in tri-culture over time was counted and normalized to the initial number of particles.

### 2.6 Cytokine profile assay

The co-cultures and tri-cultures were incubated with vehicle control or 1 μM FITC-Aβ for 96 hours. Equal volumes of conditioned media were pooled from three wells of each group for three different biological replicates with no further dilution. The media were then centrifuged at 300g for 5 minutes to settle debris, and the supernatant was preserved at -80°C. The collected samples were mailed to Eve Technologies, Calgary, AB, Canada on dry ice following the manufacturer’s instructions, where Rat Cytokine Array/Chemokine Array 27-Plex Discovery Assay was performed.

Heatmap and statistical comparisons were used to quantify and compare the amount of cytokine released in each experimental group. Standard curves were used to identify the concentration of secreted cytokines according to the relative fluorescence expression. When the standard curve was non-linear, concentration values were used instead of relative fluorescence for the cytokine. Specifically, concentration values of IP-10 (CXCL10), RANTES (CCL5) and MIP-1α (CCL3) were used due to non-linearity in the standard curve. Additionally, a logarithmic scale was used for all the values to obtain normal distribution. Hierarchical cluster analysis was performed using heatmap function in R statistical analysis package (version 4.3.1).

### 2.7 Microglial motility assessment

Following the media change on DIV 10, 1 μM FITC-Aβ was applied to mono-cultures and tri-cultures for 1 hour prior to imaging. The cells were then imaged every 5 minutes for 4 hours using a Zeiss Observer Z1 microscope with environmental control (37°C and 5% CO_2_). Cell trajectories were traced as previously described (Feng et al., 2012; Gorelik and Gautreau, 2014; Yang et al., 2019; Le et al., 2023). The “Chemotaxis and Migration Tool” (ibidi GmbH, Gräfelfing, Germany) was used to plot the migration trajectory (Zengel et al., 2011). Several motility parameters were extracted using MTrackJ plugin in Fiji/ImageJ: (i) Euclidean distance – the direct distance between the initial and final position of each cell. (ii) Migration speed – accumulated distance divided by imaging duration (i.e., 4 hours). (iii) Directionality ratio –Euclidean distance divided by the accumulated distance (Gorelik and Gautreau, 2014). For example, a highly direct trajectory results in a directionality ratio close to 1, whereas a highly random trajectory results in a ratio close to 0. To complement the analysis of the directionality ratio, we extracted the instantaneous speed across adjacent time points by dividing the Euclidean distance between the two cartesian coordinated by the duration (5 minutes). Overall, more than 150 cells (at least 50 cells per biological replicates) were assessed for each condition at every time point.

### 2.8 Statistical methods

All experiments employed at least three biological replicates with three technical replicates (culture wells or slides) for each condition. For all image analyses, a minimum of three predetermined fields was assessed per each technical replicate to account for variability within the cultures. To compare the responses of the two culture types to Aβ treatment, a two-way ANOVA was used to compare the main effects of the culture type and treatment. If the interaction was determined to be statistically significant, a post hoc Tukey’s test was utilized correcting for multiple comparison. A two-tailed unpaired parametric Welch’s t-test was carried out for comparing the microglia size between control and treatment groups. Statistical significance in all the experiments was determined when the p-value was less than 0.05.

## 3. Results

### 3.1 Aβ clearance

The number of FITC-Aβ particles in the tri-cultures was significantly decreased relative to that observed in the co-cultures (Figure 1A, B, D), where there was a monotonic decrease over the course of the experiment (Figure 1F). There was a significant reduction of particles in the tri-culture compared to the co-culture (p<0.001) (Figure 1D). Smaller particles (< 10 μm^2^) displayed a more significant reduction compared to larger aggregates (∼100 μm^2^), as shown in the histogram of particle size distribution at the end of the experiment (Figure 1E). The area under the curve (AUC) for the particle size distributions confirmed the decrease in number of particles in the tri-culture (Figure 1E inset).

**Figure 1.**
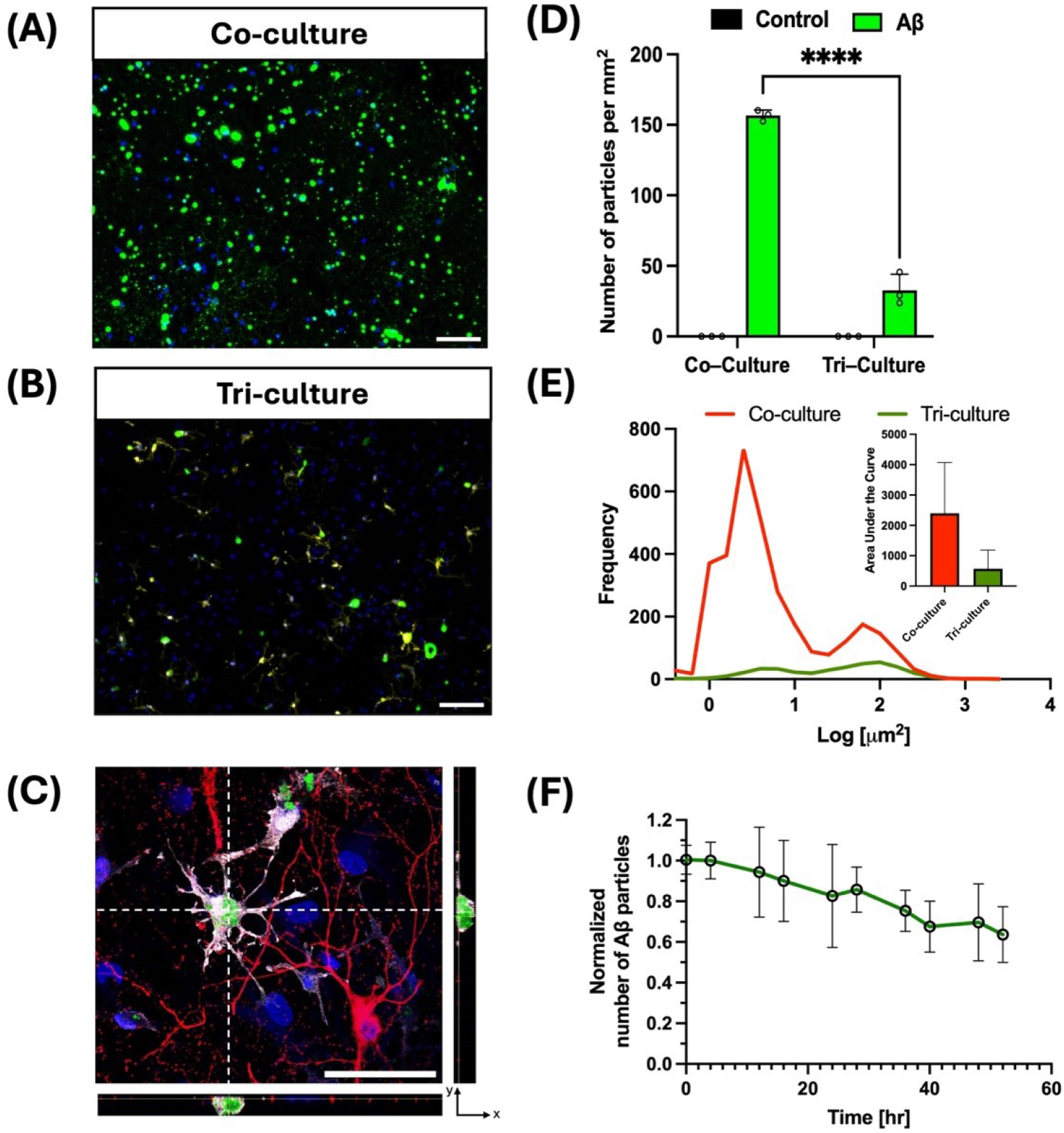
Aβ clearance analysis. Representative epifluorescence images displaying FITC-Aβ particles in (A) co-culture and (B) tri-culture after a 96-hour exposure to 1 μM FITC-Aβ initiated at DIV 10. Scale bar = 100 μm. (C) Confocal microscopy image of the tri-culture displays colocalization of FITC-Aβ with microglia. Scale bar = 50 μm. (D) Change in number of particles per FOV area and (E) size of FITC-Aβ particles in the co-culture and tri-culture (mean ± SD, n= 3 biological replicates with at least 3 wells or slides per biological replicate with 5 FOV per well/slide). (F) Number of particles (normalized to the number of particles in the initial FOV) of FITC-Aβ particles in the tri-culture was monitored every 4 hours up until 52 hours. **** p<0.0001 as determined by using two-way ANOVA with post hoc Tukey’s test.

#### 3.2 Cell morphology

Aβ exposure resulted in a significant increase in microglia size compared to the untreated control group (p<0.0001) (Figure 2A, D). In addition, the microglia size distribution widened following Aβ exposure (Figure S2). While there was a slight increase in astrocyte size due to Aβ treatment, this change did not reach statistical significance for either the co-culture or the tri-culture. Neuronal surface coverage and astrocyte size did not display a statistically-significant change upon Aβ treatment for either culture type (Figure 2 C, E). Representative astrocyte images are shown in Figure 2B.

**Figure 2.**
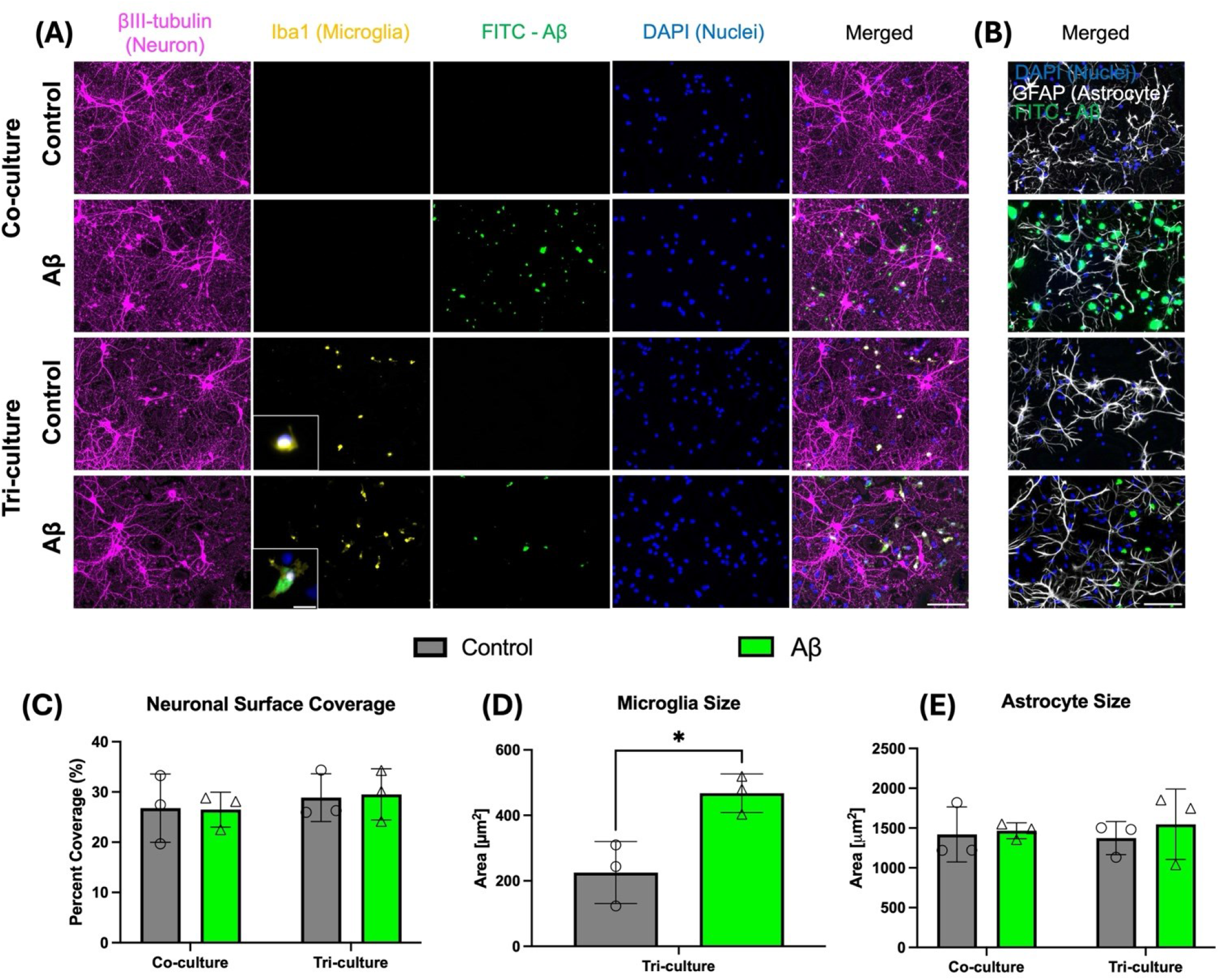
Cell morphological analysis. (A & B) Representative epifluorescence images of co-cultures and tri-cultures after a 96-hour exposure to 1 μM FITC-Aβ initiated at DIV 10. The cultures were stained to identify neurons (βIII-tubulin), microglia (Iba1), and astrocytes (GFAP). The scale bar is 100 μm in all images except for 20 μm in the inset images displaying representative single microglia. (C) Percent neuronal surface coverage. (D) Microglia size (E) Astrocyte size. These endpoints were quantified and compared using two-way ANOVA with post hoc Tukey’s test for neurons and astrocytes and Welch’s t-test for microglia size. The plots display the mean ± SD, n=3. (individual data points represents the three biological replicates with at least 3 wells or slides per biological replicate with 5 fields-of-view per well/slide). * p<0.05.

#### 3.3 Cytokine profile

The cytokine profiles of the conditioned media from the co-culture and tri-culture with/without Aβ exposure were analyzed and are presented as a concentration heatmap (Figure 3A). Among the 27 cytokines that were quantified, there was a statistically-significant change in ten cytokines for the tri-culture following Aβ exposure (Figure 3B&C), while the co-culture did not exhibit any statistically-significant changes in cytokine levels upon exposure to Aβ. Specifically, the tri-culture displayed a significantly higher concentration of the following cytokines: Cytokines that play a role in inflammation, including TNF-α (tumor necrosis factor α), IL-1α, IL-1β, IL-6, IP-10 (CXCL10), IL-10 (Figure 3B); and significantly-elevated chemokines including MIP-1α (macrophage inflammatory protein-1α), MIP-2 (macrophage inflammatory protein-2), RANTES (regulated on activation, normal T cell expressed and secreted (Figure 3C). Fractalkine (CX3CL1) concentration was decreased in the tri-culture compared to the co-culture.

**Figure 3.**
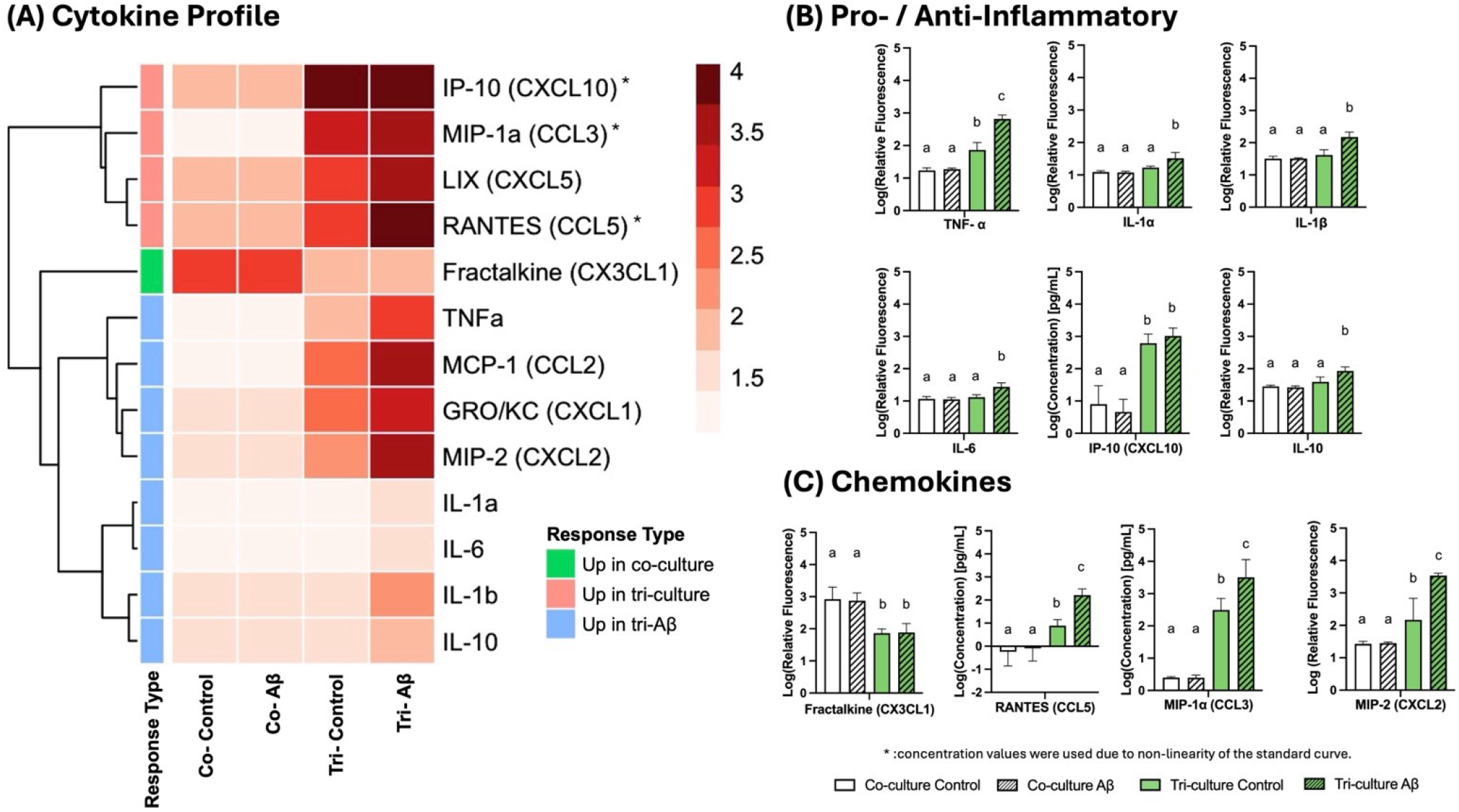
Proteomic analysis of conditioned media. (A) The cytokine heatmap displays the logarithm of relative fluorescence intensity. Only cytokines that resulted in a two-fold or higher change of the raw fluorescence intensity for any pairwise comparison between the four conditions (i.e., co-culture with/without Aβ and tri-culture with/without Aβ) are included. A full heatmap can be found in (Figure S3). Hierarchical cluster analysis revealed three major groups of cytokines that increased in the co-culture (green), increased in the tri-culture (red), and increased only in Aβ-treated tri-culture (blue). Log-scale relative fluorescence or concentration of (A) Pro- / anti-inflammatory cytokines and (B) chemokines that play a role in microglia motility. All plots display mean ± SD (n = 3 biological replicates with at least 3 wells per biological replicate pooled into one sample). Distinct letters above the bars indicate statistically-significant groups compared to the other groups, as determined with a two-way ANOVA with post hoc Tukey’s test, where p<0.05 is considered statistically-significant.

#### 3.4 Microglia motility

Microglia in the tri-culture significantly reduced their motility, e.g., Euclidian distance (Figure 4E) and directionality ratio (Figure 4G) following Aβ exposure, whereas microglia in the mono-culture did not exhibit a significant change (Figure 4A&B, Movie S1A&B). While the migratory speed was not statistically significant for microglia in the tri-culture (Figure 4F), there was a decreasing trend as a function of treatment duration (Figure S8C). Microglia in the mono-culture did not show any statistically-significant change of directionality ratio or instantaneous speed upon Aβ exposure (Figure 4G & S8B, Movie S2A&B). Individual cell-wise motility for all cells is shown as a violin plot to provide insight into motility distribution (Figure S4).

**Figure 4.**
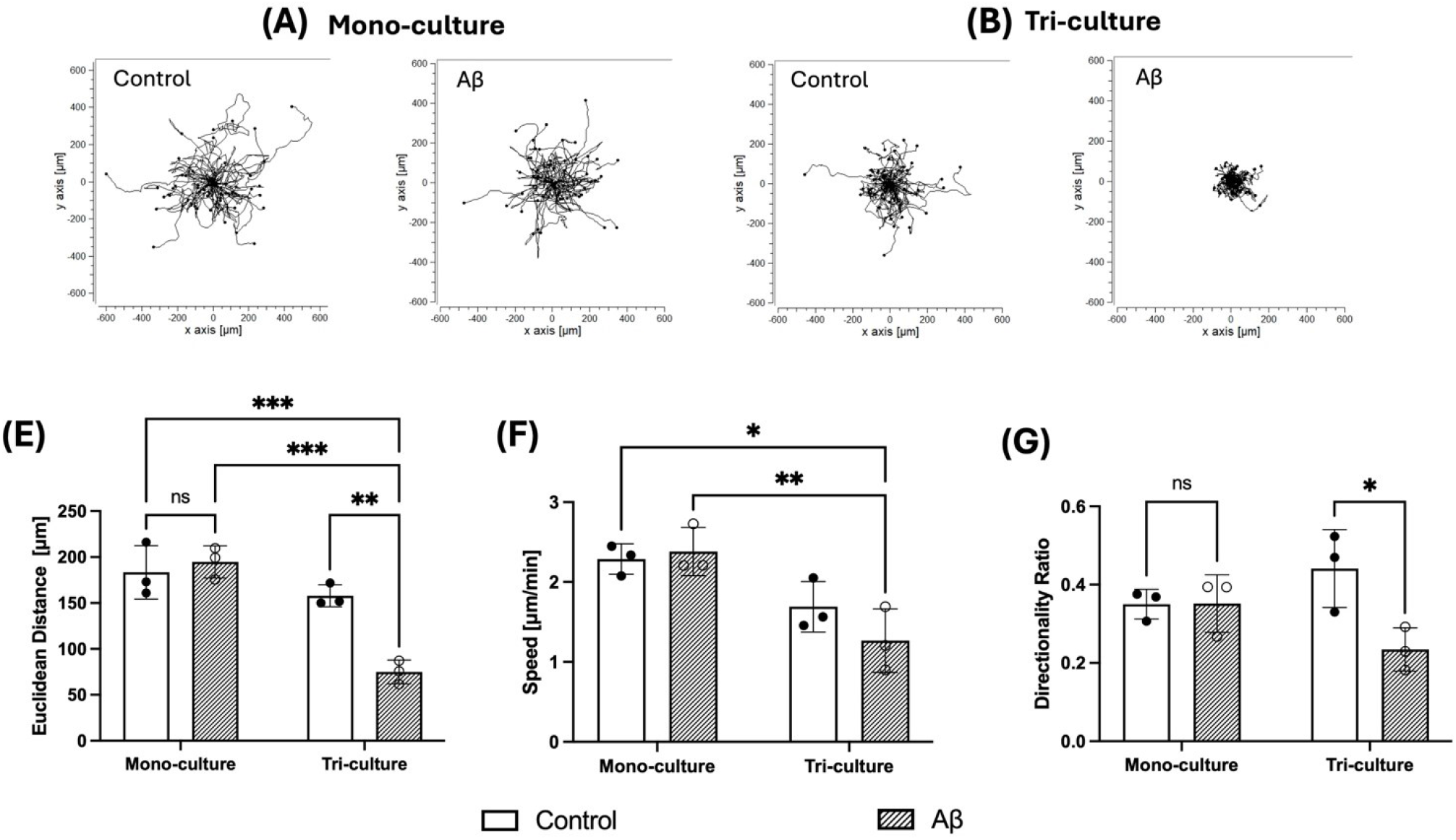
Microglia motility analysis. Mono- and tri-cultures with or without FITC-Aβ treatment were measured every 5 minutes up to 4 hours. The composite cell trajectory of microglia in (A) mono-cultures and (B) tri-cultures, with initial position of all cells translated to zero coordinate for normalization (plotted n=50 cells from one biological replicate with 5 fields of view from multiple wells). The cells were incubated with Aβ for an hour and the time-lapse images were taken for 4 hours. (E) Euclidean distance, (F) microglia speed, and (G) average directionality ratio in mono- and tri-cultures with or without FITC-Aβ treatment (mean ± SD, n=3). Two-way ANOVA with post hoc Tukey’s test was used to compare the groups, where * p<0.05, ** p<0.01, *** p<0.001 show levels of statistical significance while “ns” indicate no statistical significance.

## 4. Discussion

Here, we employed a primary rat cortical tri-culture model to characterize neural cell response to Aβ exposure with an emphasis on microglia function. The tri-culture displayed a significant reduction in fluorescent Aβ particles compared to its co-culture counterpart. This behavior is attributed to microglial phagocytosis of the fluorescently-labeled particles since *in vivo* microglia continuously survey the CNS parenchyma and clear cellular debris and/or pathogenic molecules to maintain homeostasis (Neumann et al., 2009). Furthermore, in the co-culture that lacks microglia, neuronal and astrocytic processes displayed green-fluorescent puncta, which is plausibly due to FITC-Aβ internalization by the cells and/or particle accumulation on the surface of the cells (Figure S5). In order to provide a quantitative comparison of colocalization in the co-culture and tri-culture, we analyzed pixel-wise overlap between fluorescence channel pairs (i.e., FITC-Aβ and neuron), normalized each overlap area to the total neuronal area, and compared the normalized areas using Mann-Whitney test (n=7 images) (Figure S5). There was a significant difference between the normalized overlap areas between the co-culture and tri-culture, where the normalized overlap was 6-fold higher for the co-culture suggesting that FITC-Aβ is internalized by the neurons and/or adsorbed onto neuronal surface in the absence of microglial clearance of the particles. Surprisingly, Aβ exposure did not result in any obvious changes in viability (Figure S6), morphology (Figure 2), or proteomic profile (Figure 3) in the co-culture. This may be due to shorter duration of Aβ exposure and/or decreased Aβ concentration relative to previously published studies that employed both longer durations and higher Aβ concentrations (Watson and Fan, 2005; Varghese et al., 2010; Jung et al., 2022)

On the other hand, the tri-culture displayed a much stronger response to Aβ exposure, where the presence of microglia likely plays an important role. This response is manifested at different levels. First, microglia underwent a significant morphological alteration with an increase in its cellular footprint following Aβ exposure (Figure 2D). This may at first appear to contradict what is typically observed *in vivo*, where ramified microglia become more compact with a more amoeboid morphology (Fernández-Arjona et al., 2017). However, the increase in microglia size with large lamellipodia is consistent with its *in vitro* morphology – at least due to its activation by LPS – as observed by others (abd-el-Basset and Fedoroff, 1995) and in our recent studies (Goshi et al., 2020). It is important to note that others have shown that microglia volume was influenced by the amount/size of Aβ plaques phagocytosed (Daniel Lee and Landreth, 2010), which may be partially contributing to the increased microglia size in this study. Neither the neuronal morphology or astrocytic morphology displayed significant changes upon Aβ exposure in either the co-culture or the tri-culture.

Secondly, the presence of microglia increased levels of cytokines (compared to the co-culture) with generally a further increase in the levels upon Aβ exposure (Figure 3). It is well-established that microglia secrete various cytokines upon exposure to external stimuli, including Aβ proteins, in both *in vitro* and *in vivo* studies reported by others (Hanisch, 2002; Shaftel et al., 2008; Wang et al., 2015; Decourt et al., 2017; Perez-Nievas et al., 2021). In general, the proteomic profile of the tri-culture was qualitatively similar between Aβ and LPS treatments, although the LPS treatment invoked a stronger response (i.e., higher cytokine levels) for each of the elevated cytokines (Figure S7). Among these cytokines, TNF-α (Probert, 2015), IL-1β (Mendiola and Cardona, 2018), and IL-6 (Remarque et al., 2001; Sierra et al., 2007) classically play a pro-inflammatory role. In our analyses of cytokine profiles, we observed a slight increase of TNF-α in tri-cultures compared to co-cultures, which may be due to the trauma associated with the dissociation steps (Goshi et al., 2020). Tri-cultures may be more responsive to the trauma due to early microglia activation as evidenced by the increase of IP-10 (CXCL10) via astrocytes, which is an important mediator in the initial neuroinflammatory response (Clarner et al., 2015). On the other hand, the fractalkine (CX3CL1) level, which often increases following injury due to its dissociation from neuronal membranes to recruit microglia (Sokolowski et al., 2014), was lower in the tri-culture, suggesting that the tri-culture is still in a relatively homeostatic state compared to the co-culture. Furthermore, cytokines such as MIP-2 (Sebastiani et al., 2006), MIP-1α (Rubio-Perez and Morillas-Ruiz, 2012), and IP-10 (Galimberti et al., 2006), have been reported to increase in response to Aβ particles or detected in the cerebrospinal fluid of AD patients. Intriguingly, while IL-10 typically plays a neuroprotective role and contributes to resolving inflammation (Shemer et al., 2020), we observed a significant increase following Aβ incubation. Additionally, RANTES levels were significantly higher in the tri-culture (further increased upon Aβ exposure). Others have shown that supplementing HT22 immortalized mouse hippocampal neuronal culture with RANTES reduced toxicity of Aβ exposure (Ignatov et al., 2006). Increases in IL-10 and RANTES in the tri-culture may be indicative of the dual role of microglia in leading a pro-inflammatory response along with a response to resolve the inflammatory state, where it is plausible that various microglia phenotypes exist in the tri-culture that range from neurotoxic to neuroprotective types. This diversity in microglia phenotypes is also partially reflected in the increasing distribution of microglia size (surrogate for its activation state assessed by the ramified versus ameboid morphology (Goshi et al., 2020) upon Aβ exposure (Figure S2). With the wide range of microglia phenotypes, taken together with the lack of significant morphological changes indicative of severe inflammatory response (e.g., neuronal loss and astrocyte hypertrophy) as observed in treatment with high concentrations of LPS (Goshi et al., 2020), the current model provides the opportunity to study more nuanced changes (e.g., alterations in electrophysiological activity (Goshi et al., 2023; Konstantinidis et al., 2023), where neurons should remain sufficiently viable.

In addition, some of these cytokines function are chemokines for microglia, thereby influencing their motility and migratory behavior (Nimmerjahn et al., 2005). Previous studies, involving *in vivo* imaging of microglia behavior in response to Aβ plaques in transgenic mice (Bolmont et al., 2008), along with similar *in vitro* studies with cultured microglia (Honda et al., 2001; Cho et al., 2013; Bohlen et al., 2017), indicated that the microenvironment significantly modulated microglia motility (Smolders et al., 2019). We observed that Aβ exposure significantly reduced average Euclidean distance and directionality ratio of microglia in the tri-culture. While the microglia speed in the tri-culture was significantly less compared to the mono-culture, Aβ treatment showed a tendency to decrease speed in the tri-culture; however, this difference did not reach statistical significance. It is worth noting that a statistical comparison that treats each microglia independently (hence, n=150-180) revealed a significant decrease in speed upon Aβ treatment only in the tri-culture (Figure S4). Additionally, it is important to note that while the “instantaneous speed” of microglia did not change for the mono-culture (Figure S8B) during the 4-hour live-imaging duration, microglia speed rapidly deviated from its baseline speed upon exposure to Aβ in tri-culture (Figure S8C) despite a small reduction in its magnitude. Interestingly, the reduction in directionality ratio (directional persistence), which is the ratio of the Euclidean distance to the total distance travelled, suggests that microglia motion became more random albeit within a smaller Euclidean distance, which is reminiscent of “swarming behavior” observed in macrophages and neutrophils encountering a pathogens and inflammation (Jones et al., 2014; Kienle and Lämmermann, 2016). Surprisingly, the microglia in mono-culture did not display a change in its motility upon Aβ exposure. Microglia closely interact with other CNS cells via soluble cues and cell-cell contact (Angiari et al., 2022), therefore, this observation suggests that signals from neurons and astrocytes in the tri-culture play an important role in shaping microglia motility (Garland et al., 2022). One such example is the reduced CX3CL1/fractalkine levels in the tri-culture compared to the co-culture, indicating that majority of membrane-bound CX3CL1 in tri-cultures bonds and interacts with microglia plausibly decreasing the average microglia speed due to microglia-neuronal interactions (Goshi et al., 2020). Other studies reported that IP-10 (CXCL10) (Rappert et al., 2004), RANTES (CCL5) (Haruwaka et al., 2019), and MIP-1α (CCL3) (Wang et al., 2008; Zhao et al., 2018) also influenced microglial motility via interactions between neurons and astrocytes. CCL5 and CCL3 are the ligands of CCR5 receptor on microglia, which is a prominent chemokine receptor for migratory behavior (Carbonell et al., 2005). Therefore, it is reasonable to at least partially attribute the changes in microglia motility in the tri-culture to the elevated levels of these cytokines only in the tri-culture with an augmented effect for Aβ treatment group (Figure 3), where the influence of chemokines on microglia motility has been shown by others (Rappert et al., 2004; Dou et al., 2012). In addition, external cues including culture surface topography (Pires et al., 2015) and neuronal activity (Badimon et al., 2020; Nebeling et al., 2023) provide additional signals for microglia migration. Collectively, it is evident that the presence of neurons and astrocytes significantly influenced microglia behavior.

## Conclusion

Here, we have described the utility of the tri-culture model for studying microglia function and its potential use in investigating microglial response to Aβ, an important contributor to the pathogenesis of AD. Future studies should include transcriptomic analysis of microglia phenotypes and influence of induced inflammation on Aβ clearance. Several studies allude to the influence of Aβ proteins of different physical properties (soluble vs plaque-like) on neuronal activity and the influence on neuronal activity and microglial synaptic pruning and clearance. Therefore, future studies should also focus on studying the interplay of Aβ peptide forms (e.g., soluble monomeric and oligomeric, aggregates, plaques), electrophysiology, and microglial clearance.

## Author Contributions

H.K. and E.S. designed the experiments and performed the primary interpretation of the results. H.K. performed cell culture, imaging, and data analysis, and wrote the original manuscript. B.L. and K.Z. contributed to the microglia motility imaging and data analysis. N.G., A.C.G, M.Z., and P.J.L contributed to the interpretation of the experimental results and edited the manuscript. All authors have read and agreed to the published version of the manuscript.

## Acknowledgments

The authors are grateful to Dr. Johannes Hell and Dr. Sarah Rogue at UC Davis for their help on characterizing FITC-Aβ particles with silver staining and immunoblotting. In addition, we thank Arushi Patel for her help with microglia motility analysis and astrocyte area characterization, and Marion Hardy for helping with hierarchical clustering of cytokine data.

## Supplementary Material

Supplementary Material document includes details on Aβ particle analysis, cytotoxicity analysis due to Aβ treatment, cytokine panel heatmap showing co-culture and tri-culture responses to Aβ and LPS, microglia population statistics, and description of the supplemental microglia motility videos.

## Data Availability Statement

The datasets generated during and/or analyzed during the current study are available from the corresponding author on reasonable request.

