## Supplementary Material for "Rat primary cortical cell tri-culture to study effects of amyloid-beta on microglia function"

### 1 Supplementary Data

#### 1.1 FITC-A$\beta_{1-42}$ characterization

FITC-Aβ preparations were first characterized by silver nitrate staining. Briefly, FITC-Aβ solution was mixed with 1.5x sodium dodecyl-sulfate (SDS) sample buffer (4.5% SDS, 1.7% (or 140 mM) Tris, 14% sucrose, 0.006% bromophenol blue) and added to each well of 4-20% gradient precast polyacrylamide gel (Bio-Rad) for electrophoretic separation. A BioRad Dual Extra sizing ladder (2 kDa to 250 kDa) was included on the gels to evaluate the molecular weight of the FITC-Aβ preparations.

On a separate gel, immunoblotting was conducted to confirm the presence of Aβ peptides (Figure S1). Briefly, the FITC-Aβ peptides were added to 2.5% β-mercaptoethanol solution and incubated at 95°C in a heating block for 5 minutes to separate protein subunit before adding them to individual wells of a polymerized 15% acrylamide gel. Electrophoresis was performed at 100 V for an hour. After the proteins were electro-transferred to polyvinylidene difluoride (PVDF) membrane (12 mA for 45 min), the blot was blocked for 1 hour with 5% milk in TBS (TBS-milk) and incubated for 1 hour with anti-β-amyloid, 1-16 antibody (diluted 1:2500, BioLegend, RRID: AB_2715854). The gel was washed three times with Tween 20 (0.1%) in TBS (TBS-T) for 10 minutes each at room temperature, followed by secondary antibody incubation that contained Peroxidase AffiniPure goat anti-mouse IgG (1:10,000, Jackson Immuno Research, RRID: AB_2338503) for 1 hour at room temperature. Finally, the gel was washed twice in TBS-T followed by a single TBS wash for 10 minutes at room temperature and was treated with Immobilon Classico Western HRP substrate (Millipore) for 3 minutes. The film was imaged using a film processor (SRX 101A Konica Minolta) to visualize the bands.

#### 1.2 Lactate dehydrogenase assay

The cell cytotoxicity of the FITC-Aβ preparation at varying doses on the co-culture and tri-culture was assessed using CyQUANT™ LDH Cytotoxicity Assay (Invitrogen) per the manufacturer's instructions (Figure S6). Prior to using the assay, the conditioned media was collected, centrifuged at 300g for 5 minutes, and the supernatant was collected to eliminate the introduction of dead cells. The values were normalized to the median of the control (vehicle) group. One-way ANOVA was used and comparisons to the control group was conducted via post hoc Dunnett's test.

#### 1.3 Cytokine profile of conditioned media from co-culture and tri-culture treated with Aβ or LPS

The co-cultures and tri-cultures were incubated with vehicle (control) or 5 μg/mL LPS for 96 hours. Equal volumes of conditioned media were pooled from the three wells of each condition for each biological replicate with no further dilution. For 1 μM FITC-Aβ, three biological replicates were included. The media was then centrifuged at 300g for 5 minutes to settle debris, and the supernatant was preserved at -80°C. The collected samples were mailed to Eve Technologies, Calgary, AB, Canada on dry ice following the manufacturer's instructions, where Rat Cytokine Array/Chemokine Array 27-Plex Discovery Assay was performed (Figure S7).

#### 1.4 Colocalization of the FITC-Aβ to neurons in co-culture and tri-culture

Seven images (at 100x magnification) stained for βIII-tubulin (neurons) and Iba1 (microglia) from one biological experiment were chosen to evaluate the co-localized (overlap) area between β-III tubulin (neurons) and FITC-Aβ in co- and tri-cultures (Figure S5). Images were processed using FIJI/ImageJ BIOP JACoP plugin. Auto threshold method 'Li' was used for all images. The ratio of overlapped area was calculated by normalizing overlap area values to the neuronal cell surface coverage area of each image.

### 2. Supplementary Figures and Tables

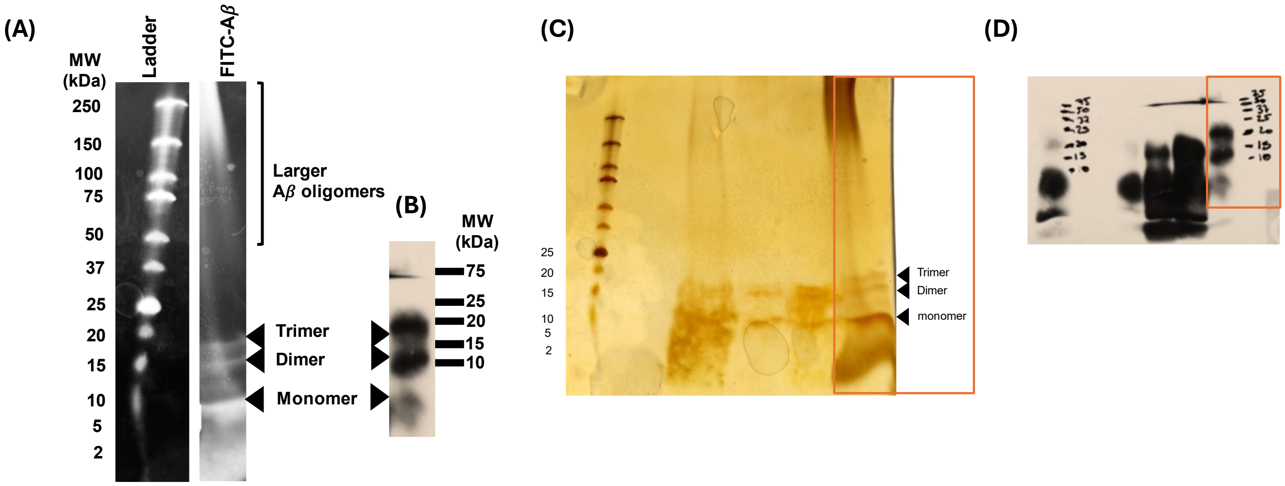


**Figure S1.** Gel electrophoresis to characterize molecular weights of FITC-Aβ peptides in solutions used for the experiments (A) silver staining for non-specific protein visualization, (B) immunoblotting for specific Aβ visualization, and (C&D) original images of silver staining and immunoblotting (Orange box highlights the lane of the treatment).

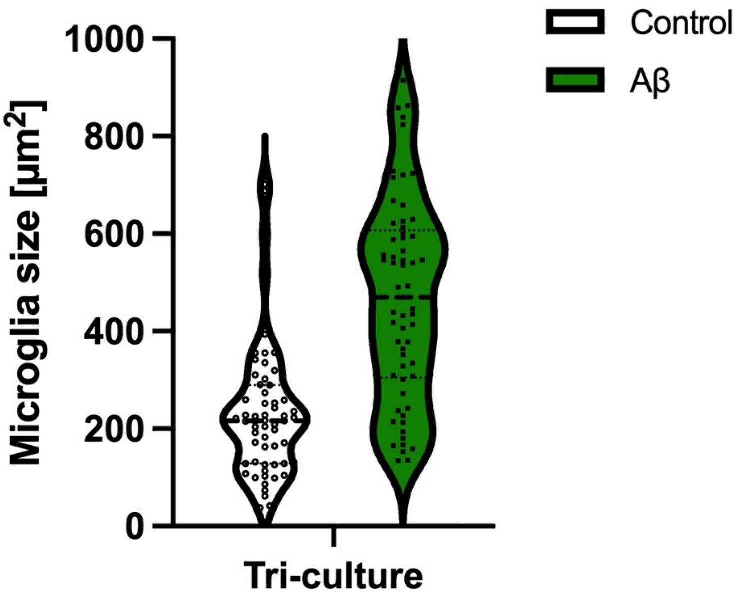


**Figure S2.** Microglial size distribution with or without Aβ treatment, where each data point is an individual microglia.

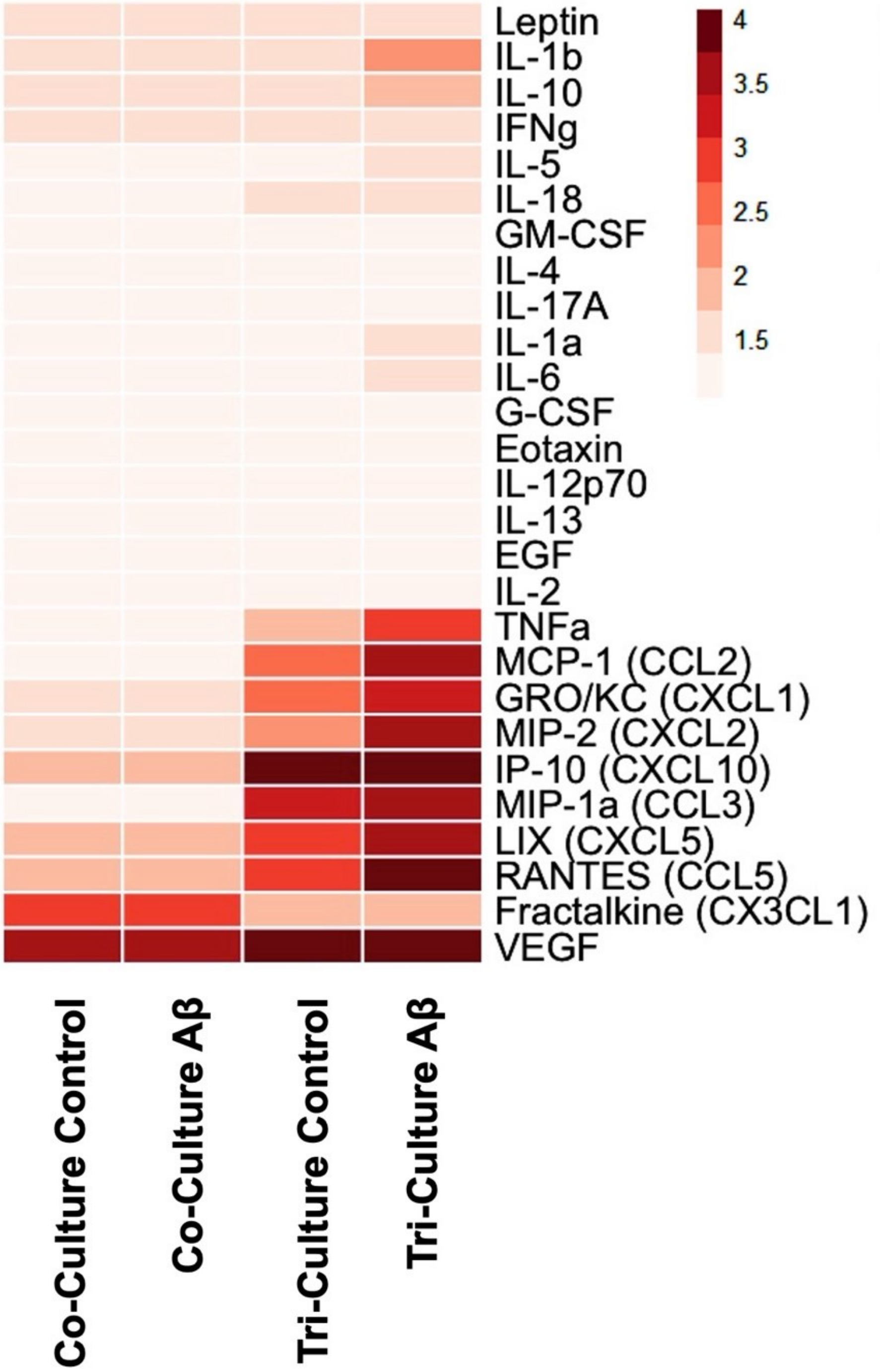


**Figure S3.** Heatmap of entire set of cytokine levels (logarithm of relative fluorescence intensity) shown in Figure 3A.

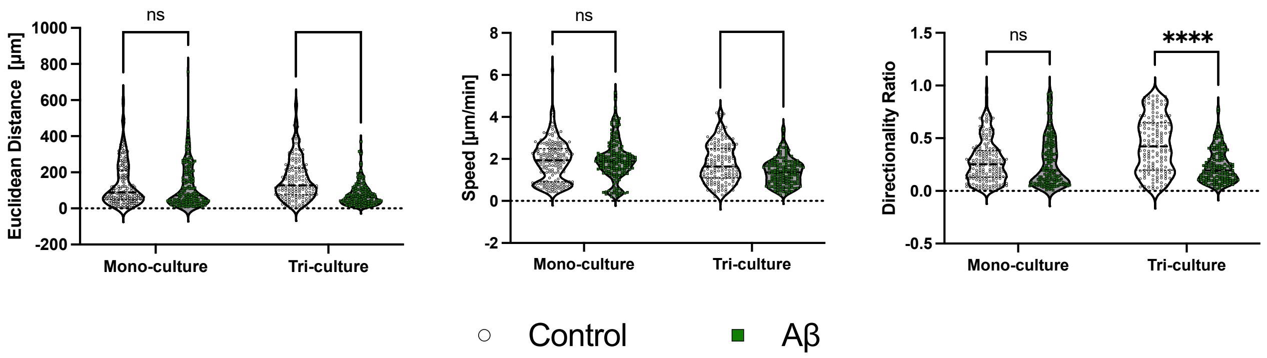


**Figure S4.** Violin plot of all the cells (n=150-180) analyzed to extract motility parameters (i.e., Euclidean distance, speed, directionality ratio). A multiple comparison including all the individual cell types was conducted via a two-Away ANOVA with post hoc Tukey test. *** p<0.001, **** p<0.0001, ns: not significant.

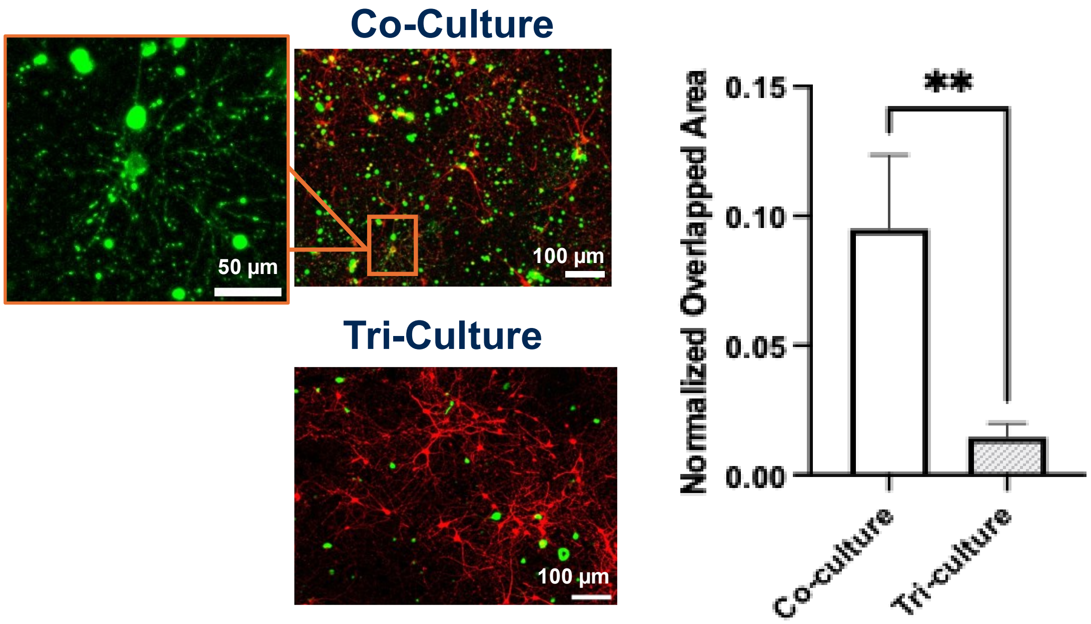


**Figure S5.** Co-localization of FITC-Aβ particles to neuronal cells (β-III tubulin positive) in the co-culture and tri-culture. (7 pooled images from one biological experiment with at least 3 wells with 5 field of view per well). The overlap area is normalized to the neuronal surface coverage for each image. Mann-Whitney test was conducted. **p<0.01, ns: not significant.

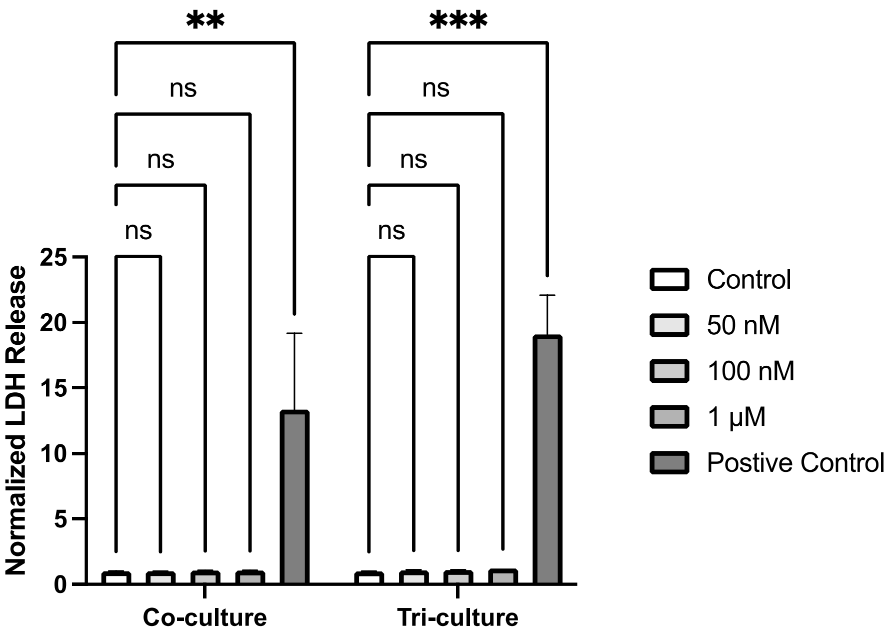


**Figure S6.** Dose-response cell toxicity of cultures treated with FITC-Aβ for 96 h initiated at DIV 10. Comparing the control condition in co- and tri-cultures to varying FITC-Aβ concentrations, including the Triton X-100 treated positive control. The LDH release (surrogate for cell toxicity) for each FITC-Aβ concentration was compared to the control via Dunnett's multiple comparison statistics. (n=2 biological replicates with at least 3 wells per biological replicate). ** $p<0.01$, *** $p<0.001$, ns: not significant.

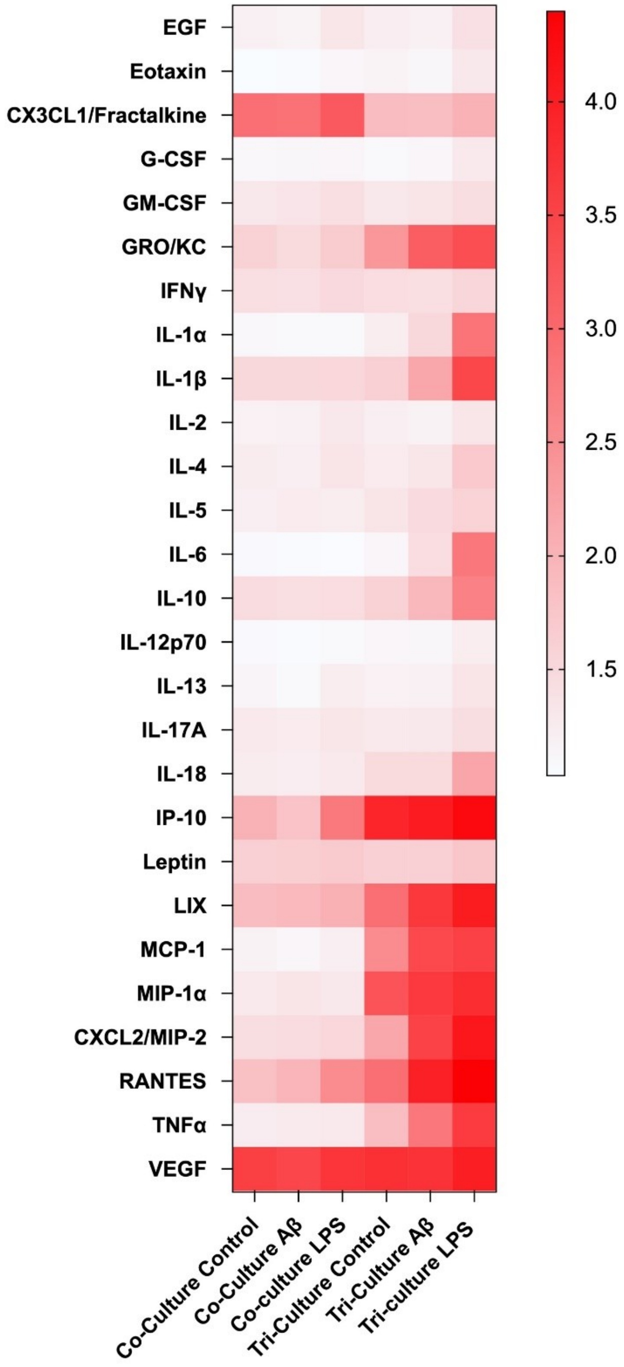


**Figure S7.** Heatmap of cytokine levels (logarithm of relative fluorescence intensity) of conditioned media from the co-cultures and tri-cultures treated with or without FITC-Aβ and LPS (n=3 biological replicate for FITC-Aβ and n=1 biological replicate for LPS with at least 3 wells pooled into one sample).

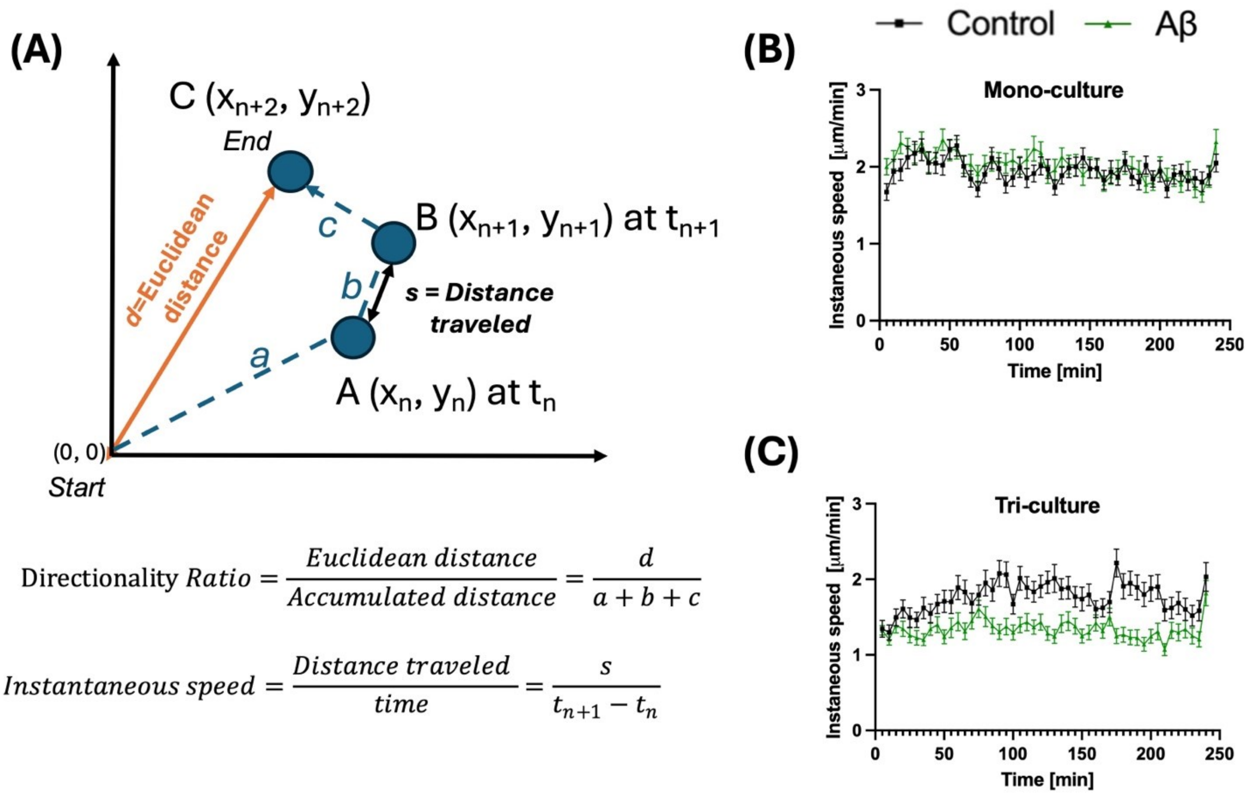


**Figure S8.** (A) Graphical and mathematical descriptions of the motility parameters. Instantaneous speed of each cell (n=150-180 cells total from 3 biological replicates with at least 5 fields of view from multiple wells) in (B) mono-culture and (C) tri-culture with or without FITC-Aβ were measured every 5 minutes up to 4 hours.

**Movie S1A.** Time-lapse movie for tri-culture in vehicle control (Image size: 2365 µm x 1936 µm)

**Movie S1B.** Time-lapse movie for tri-culture in FITC-Aβ (Image size: 2365 µm x 1936 µm)

**Movie S2A**. Time-lapse movie for mono-culture in vehicle control (Image size: 1183 µm x 968 µm)

**Movie S2B**. Time-lapse movie for mono-culture in FITC-Aβ (Image size: 1183 µm x 968 µm)
